# Vps74 connects the Golgi apparatus and telomeres in *Saccharomyces cerevisiae*

**DOI:** 10.1101/269456

**Authors:** Joana Rodrigues, Peter Banks, David Lydall

**Author notes:** Corresponding author: David Lydall, Faculty of Medical Sciences, Framlington Place, Institute for Cell and Molecular Biosciences, Cookson Building (M2.022), Newcastle University, Newcastle, NE2 4HH, United Kingdom.

## Abstract

In mammalian cell culture, the Golgi apparatus fragment upon DNA damage. GOLPH3, a Golgi component, is a phosphorylation target of DNA-PK after DNA damage and contributes to Golgi fragmentation. The function of the yeast (*Saccharomyces cerevisiae*) ortholog of GOLPH3, Vps74, in the DNA damage response has been little studied, although genome-wide screens suggested a role at telomeres. In this study we investigated the role of Vps74 at telomeres and in the DNA damage response. We show that Vps74 decreases the fitness of telomere defective *cdc13-1* cells and contributes to the fitness of *yku70Δ* cells. Importantly, loss of Vps74 in *yku70Δ* cells exacerbates the temperature dependent growth defects of these cells in a Chk1 and Mec1-dependent manner. Furthermore, Exo1 reduces fitness of *vps74Δ yku70Δ* cells suggesting that ssDNA contributes to the fitness defects of *vps74Δ yku70Δ* cells. Systematic genetic interaction analysis of *vps74Δ, yku70Δ* and *yku70Δ vps74Δ* cells suggests that *vps74Δ* causes a milder but similar defect to that seen in *yku70Δ* cells. *vps74Δ* cells have slightly shorter telomeres and loss of *VPS74* in *yku70Δ* or *mre11Δ* cells further shortens the telomeres of these cells. Interestingly, loss of Vps74 leads to increased levels of Stn1, a partner of Cdc13 in the CST telomere capping complex. Overexpression of Stn1 was previously shown to cause telomere shortening, suppression of *cdc13-1* and enhancement of *yku70Δ* growth defects, suggesting that increased levels of Stn1 may be the route by which Vps74 affects telomere function. These results establish Vps74 as a novel regulator of telomere biology.

## INTRODUCTION

The Golgi apparatus is found in all eukaryotes, functioning in the maturation of proteins and lipids destined for the cell surface or other internal compartments (Glick and Nakano 2009; Potelle et al. 2015). Somewhat surprisingly, in mammalian cells, the Golgi responds to DNA damage (Farber-Katz et al. 2014). Golgi becomes fragmented after camptothecin (CPT)-induced DNA damage and Golgi fragmentation persists long after DNA lesions are repaired. The fragmentation in response to DNA damage was dependent on GOLPH3, which was phosphorylated by DNA-PK (a DNA damage protein kinase). GOLPH3 phosphorylation increased its interaction with MYO18A, a myosin that links Golgi membranes to the cytoskeleton (Dippold et al. 2009).

*GOLPH3* is highly conserved among eukaryotes and *VPS74* is its budding yeast (*Saccharomyces cerevisiae*) orthologue. Vps74 is reported to be important for the localisation of glycosyltransferases to the Golgi apparatus and to activate Sac1, a phosphoinositide phosphatase membrane protein, in Golgi (Schmitz et al. 2008; Wood et al. 2012). Glycosyltransferases are responsible for protein glycosylation, where sugar (glycan) chains are attached to proteins, contributing to the correct folding and function of these proteins (Shental-Bechor and Levy 2008; Xu and Ng 2015). Sac1 regulates the levels of phosphatidylinositol 4-phosphates (PtdIns4P) which are lipids known to promote protein trafficking in Golgi (Strahl and Thorner 2007). Phosphatidylinositols can also affect nuclear mRNA export, with a decrease in InsP6 (whose precursor is PtdIns4P) levels, leading to an accumulation of polyadenylated mRNA in the nucleus (Wera et al. 2001). Interestingly, large-scale surveys suggested that *VPS74* affected fitness of telomere defective *yku70Δ* and *cdc13-1* cells in opposite directions (Addinall et al. 2011). Here we carefully examined the role of *VPS74* in telomere defective cells. Low and high throughput data suggest that Vps74 and Yku70 work in parallel pathways to contribute to telomere capping and that Vps74 may affect telomere function by affecting levels of the critical telomere capping protein Stn1.

## MATERIALS AND METHODS

### Yeast strains

Standard procedures for yeast culture, mating and tetrad dissection were followed (Adams et al. 1997). Unless otherwise stated, all experiments were performed using *Saccharomyces cerevisiae* W303 (*RAD5*) strains as listed in Table S1. Gene disruptions were made in diploids using one step PCR to insert a kanMX or natMX cassettes into the genome (Goldstein and McCusker 1999). Gene disruptions were confirmed by PCR. Oligonucleotide sequences are available upon request.

### Yeast growth assays

A pool of colonies (>10) were grown until saturation overnight at 23°C (*cdc13-1* strains) or 30°C (other strains) in 2 ml of liquid YEPD (supplemented with adenine) or –LEU medium (for strains carrying plasmids). 5 or 7-fold serial dilutions in water were spotted onto YEPD or –LEU plates using a replica plating device. Plates were incubated for 2 or 3 days at the appropriate temperatures before being photographed. Unless stated otherwise, a single plate per temperature is shown for each figure (round plates fit between 8 and 16 strains while rectangular plates fit between 16 and 32 strains). For passage tests, single colonies (from germination plates) were streaked onto a YEPD plate and then several colonies (>10) from this plate were restruck on a new YEPD plate for each passage. Cells were grown for two days at 30°C. ImageJ (http://imagej.nih.gov/ij/) quantification was performed as outlined at http://lukemiller.org/index.php/2010/11/analyzing-gels-and-western-blots-with-image-j/.

### Analysis of telomere structure

Southern blot analysis was used to assess telomere length and performed as previously described (Dewar and Lydall 2010). Genomic DNA was extracted, digested with XhoI and then run overnight on a 1% agarose gel at 1V/cm. Southern transfer was performed using a Biorad Vacuum Blotter according to manufacturer’s indications. Y’+TG probe labelling and Southern detection were made according to the DIG High Prime DNA Labelling and Detection Starter Kit II (Roche) manufacturer’s instructions. The probe is approximately 1 kb with ~880bp of Y’ and 120bp of TG repeats and was released from pDL1574 using XhoI and BamHI.

### QFA

Query strains were created as described in Table S1 in using a PCR based lithium acetate method followed by crossing and random spore analysis. SGA (synthetic genetic array) was performed as previously described, crossing *vps74Δ*, *yku70Δ*, *vps74Δ yku70Δ* with part of the genome-wide single gene deletion knock out collection (Table S2) (Tong et al. 2001; Tong and Boone 2006). For QFA, strains from the final SGA plates were inoculated robotically into 200 μl liquid media in 96-well plates and grown for 2 days at 20°C without shaking, as previously described (Dubarry et al. 2015). After resuspension saturated cultures were spotted onto solid agar plates and were incubated and imaged as described before (Addinall et al. 2011; Dubarry et al. 2015). In order to measure fitnesses of the various query strains at high temperatures, a total of eight replicates of QFA were performed at both 36°C and 37°C. Fitness and genetic interaction strength estimates were performed as described before (Dubarry et al. 2015).

### Western blots

Protein extracts were prepared by trichloroacetic acid (TCA) precipitation (Ngo and Lydall 2015). Briefly, cells were resuspended in 10% TCA and mechanically broken using glass beads. Protein suspensions in Laemmli buffer were boiled for 3 min, spun down for 10 min and the supernatant were loaded onto 4-15% Mini-PROTEAN TGX Gels (Bio-Rad). The proteins were transferred to a nitrocellulose membrane (GE Healthcare) and probed with anti-Myc (Abcam ab32), anti-tubulin antibodies (from Keith Gull, Oxford University) and anti-Rad53 (Abcam ab104232).

### Data availability

All strains and materials are available upon request. File S1 contains the supplementary Figures S1-S7. File S2 contains the raw data from the QFA screens performed in this study. Table S1 contains the list of strains used in the study. Table S2 contains the genes analysed in the screens. Table S3 describes the plasmids used in this study.

## RESULTS

### Vps74 is important for telomere biology

Yeast genome-wide screens in the S288C genetic background suggested that *VPS74* is involved in telomere biology (Addinall et al. 2011). Deletion of *VPS74* weakly suppressed *cdc13-1* fitness defects at 27°C and strongly enhanced *yku70Δ* fitness defects at 37°C (Addinall et al. 2011). In order to clarify the role of *VPS74* at telomeres, *VPS74* was deleted in *cdc13-1* and *yku70Δ* telomere defective cells in the W303 genetic background and the fitness of the double mutants was carefully assessed by spot test.

In agreement with the high-throughput data, *vps74Δ cdc13-1* cells grow better at 27°C than *cdc13-1* cells, showing that Vps74 reduces the fitness of *cdc13-1* cells (Figure 1A). On the other hand, *vps74Δ yku70Δ* cells are significantly less fit than *yku70Δ* or *vps74Δ* cells at 36°C, showing that Vps74 is important for the fitness of *yku70Δ* cells (Figure 1B). We note that *vps74Δ* cells grew poorly at 23°C (Figure 1). We conclude that Vps74 slightly decreases the fitness of *cdc13-1* cells but increases the fitness of *yku70Δ* cells.

**Figure 1.**
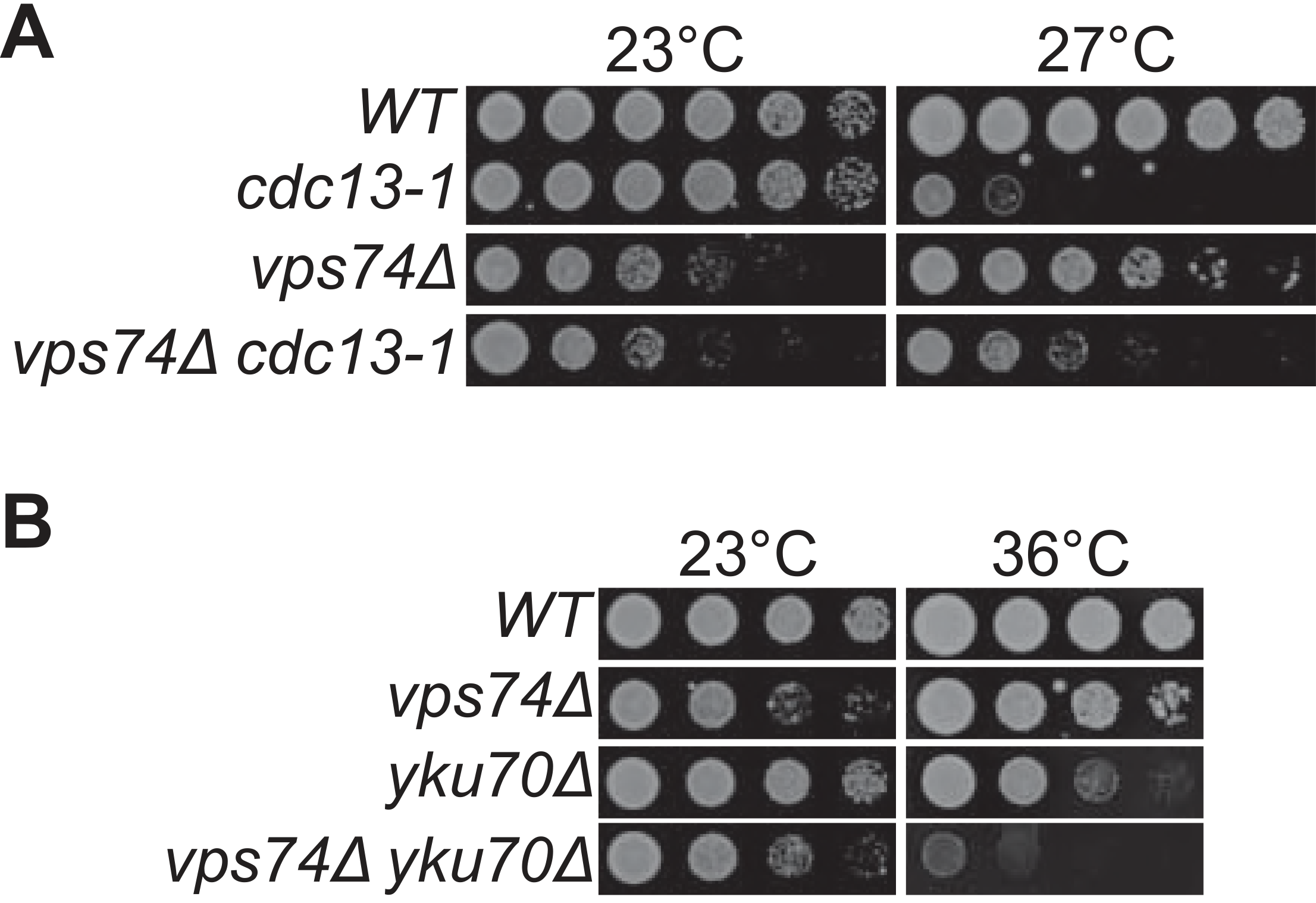
*VPS74* affects the fitness of telomere defective cells. **(A and B)** Serial dilutions of saturated overnight cultures, grown at 23°C, were spotted onto YEPD plates and incubated for 2 days at the indicated temperatures. All strains are in the W303 genetic background and at each temperature were grown on a single plate but images have been cut and pasted to allow better comparisons.

### The DDR checkpoint is activated in *vps74Δ yku70Δ* cells

To better understand the molecular nature of the defect in *vps74Δ yku70Δ* cells, we measured genetic interactions with gene deletions affecting the DNA damage response using *vps74Δ* and *yku70Δ* single mutants as controls. We tested *CHK1, MEC1, EXO1* and *MRE11* since these all affect fitness of *yku70Δ* cells (Maringele and Lydall 2002). Chk1 and Mec1 were shown to be important for cell cycle arrest in *yku70Δ* cells and Exo1 is the major exonuclease responsible for telomeric DNA resection in these mutants (Maringele and Lydall 2002). Mre11, part of the MRX complex, is important for *yku70Δ* cell fitness and simultaneous loss of Mre11 and Yku70 leads to extensive telomere rearrangements (Maringele and Lydall 2004). *chk1Δ* strongly suppressed *yku70Δ vps74Δ* fitness, and *yku70Δ* fitness defects as expected, but did not affect *vps74Δ* fitness at 38°C (Figure 2A). *mec1Δ* (*sml1Δ*) suppressed *yku70Δ vps74Δ, yku70Δ* and *vps74Δ* fitness defects, at 35°C and at 38°C, respectively (Figure 2B). *exo1Δ* strongly suppressed *vps74Δ yku70Δ* and *yku70Δ* fitness defects, allowing these cells to grow at 38°C (Figure 2C). *vps74Δ exo1Δ* cells grew similarly to *vps74Δ* cells at 38°C (Figure S1A and Figure 2C). In contrast, *mre11Δ* strongly enhanced *vps74Δ* fitness defects at 23°C and 37°C (Figure 2D).

**Figure 2.**
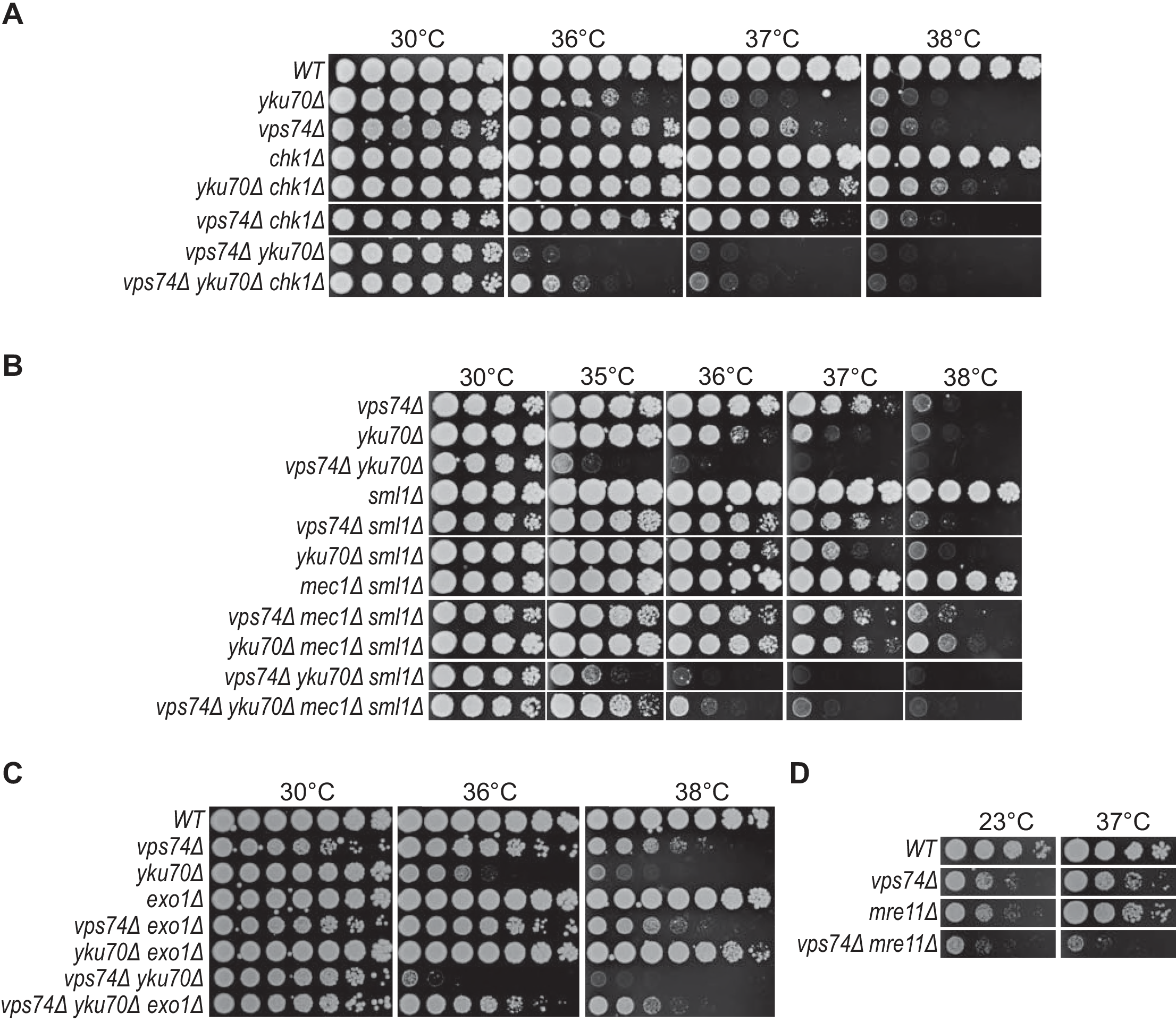
*CHK1, MEC1, EXO1* and *MRE11* affect the fitness of *vps74Δ yku70Δ* and *vps74Δ* cells at high temperatures. Spot test assays as described in Figure 1.

Given that Exo1 decreases the fitness of *vps74Δ yku70Δ* cells when compared to *yku70Δ* cells, we hypothesised that loss of *VPS74* leads to increased telomeric ssDNA in *yku70Δ* cells. To test this, telomeric ssDNA levels in *vps74Δ* and *vps74Δ yku70Δ* cells was measured by in-gel assay after 4h growth at 36°C. However, loss of *VPS74* did not strongly affect the levels of telomeric ssDNA in *WT* or *yku70Δ* cells (Figure S1C-E). Interestingly, *vps74Δ* decreased the ssDNA levels of *cdc13-1* cells as measured by both In-gel assay and quantitative amplification of ssDNA (QAOS) (Figure S1C-E). This decrease in ssDNA is in agreement with the suppression of *cdc13-1* fitness defects by *vps74Δ* (Figure 1A), since increased levels of ssDNA in *cdc13-1* cells were shown to be responsible for the poor fitness of these cells (Maringele and Lydall 2002). Together these data suggest that Vps74 functions with Yku70 and Mre11 to help cap telomeres and protect from DDR pathways.

### Loss of Vps74 leads to telomere shortening

Since *vps74Δ mre11Δ* cells have extensive fitness defects at all temperatures we wondered if *vps74Δ,* like *yku70Δ,* when combined with *mre11Δ,* leads to progressive telomere attrition and survivor appearance (Maringele and Lydall 2004). To test this, we passaged *vps74Δ mre11Δ* cells and analysed their colony size and telomere lengths. We observed that the fitness of *vps74Δ mre11Δ* cells improves with passage (Figure 3A, B). Interestingly, *vps74Δ* cells show slightly short telomeres, consistent with a role for Vps74 in telomere capping, but the effect is much less than either Yku70 or Mre11 (Figure 3C). Telomeres of *vps74Δ mre11Δ* were slightly shorter than telomeres of *mre11Δ* cells (Figure 3C). Furthermore, telomeres of *vps74Δ mre11Δ* cells slightly lengthen with passage but do not show the major telomere rearrangements seen in telomerase deficient survivors (Maringele and Lydall 2004). These data show that Vps74 has a minor role affecting telomere length, independent of Mre11 and Yku70.

**Figure 3.**
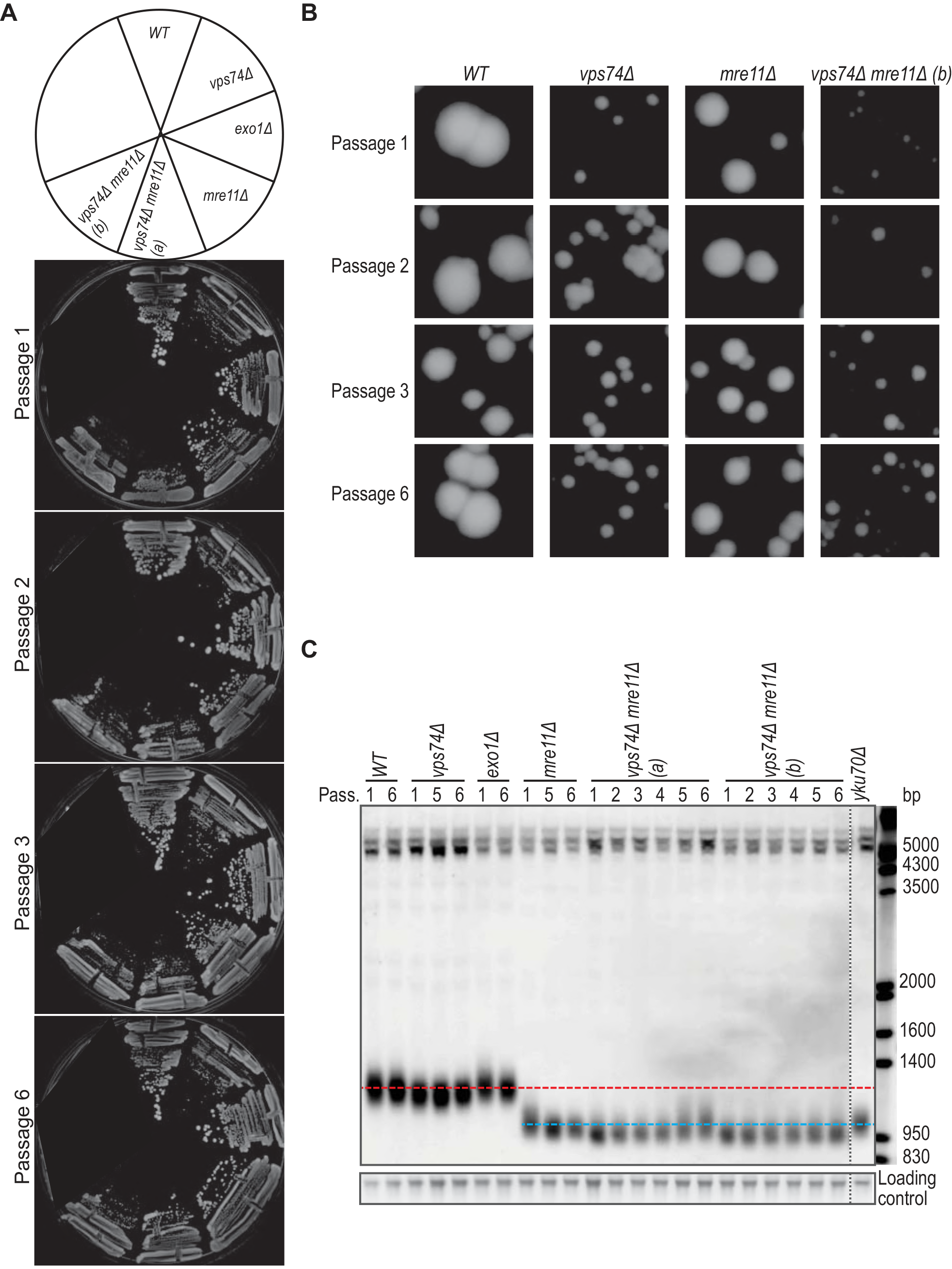
Loss of *VPS74* leads to telomere shortening in *WT* and *mre11Δ* cells. **(A)** Passage tests performed at 30°C. Cells were allowed to grow for 2 days before pictures were taken and cells passaged. **(B)** Zoom in of the colonies in A. **(C)** Cells from the plates in A were inoculated in liquid YEPD, grown until saturation and DNA was extracted. The DNA was analysed by Southern blot with a telomere probe (Y’+TG). Horizontal red line represents the *WT* telomere length and the blue line is roughly the telomere length of *mre11Δ* cells. Vertical dashed line indicates where the gel picture was cut for presentation purposes.

### Large-scale studies suggest that Vps74 has similar, but parallel, functions to Yku70

To systematically explore the role of Vps74 in telomere function, a medium scale quantitative fitness analysis was conducted to compare genetic interactions of 358 gene deletions in *vps74Δ*, *yku70Δ* and *vps74Δ yku70Δ* cells (Addinall et al. 2011). The 358 gene deletions were chosen because they affect DNA damage responses, intracellular protein transport, protein localization, protein maturation, regulation of phosphatidylinositol dephosphorylation, retrograde vesicle-mediated transport (Golgi to ER), telomere maintenance, endoplasmic reticulum unfolded protein response and others (Table S2). Before embarking on these screens we first confirmed that *vps74Δ* and *yku70Δ* are synthetically sick in the S288C genetic background, used for high-throughput screens (Figure S2A).

The fitness of the double and triple mutants was then assessed by QFA (Figure 4). In part to assess the quality the data we highlight gene deletions known to interact with telomere defective strains *(MRE11, EST1, CHK1, EST2, RAD50, DDC1, EXO1, RIF1, RAD9, RAD17, RAD24, TEL1, RRM3, NMD2, and YKU80).* Interestingly, although *vps74Δ* cells are not as temperature sensitive as *yku70Δ* cells (or *vps74Δ yku70Δ* cells), it is possible to see that the relative position of the highlighted genes is very similar in *vps74Δ*, *yku70Δ*, *vps74Δ yku70Δ* (Figure 4A-C). For instance, *exo1Δ* and *chk1Δ* are found as suppressors in all screens, while *nmd2Δ* and *rad50Δ* are enhancers in all screens. This pattern across the screens suggests that *VPS74* deletion causes fitness defects that have a similar origin to those observed in *yku70Δ* cells. We conclude that Vps74 and Yku70 have similar functions in the maintenance of the genome/telomere integrity, however Vps74 contribution is more minor.

**Figure 4.**
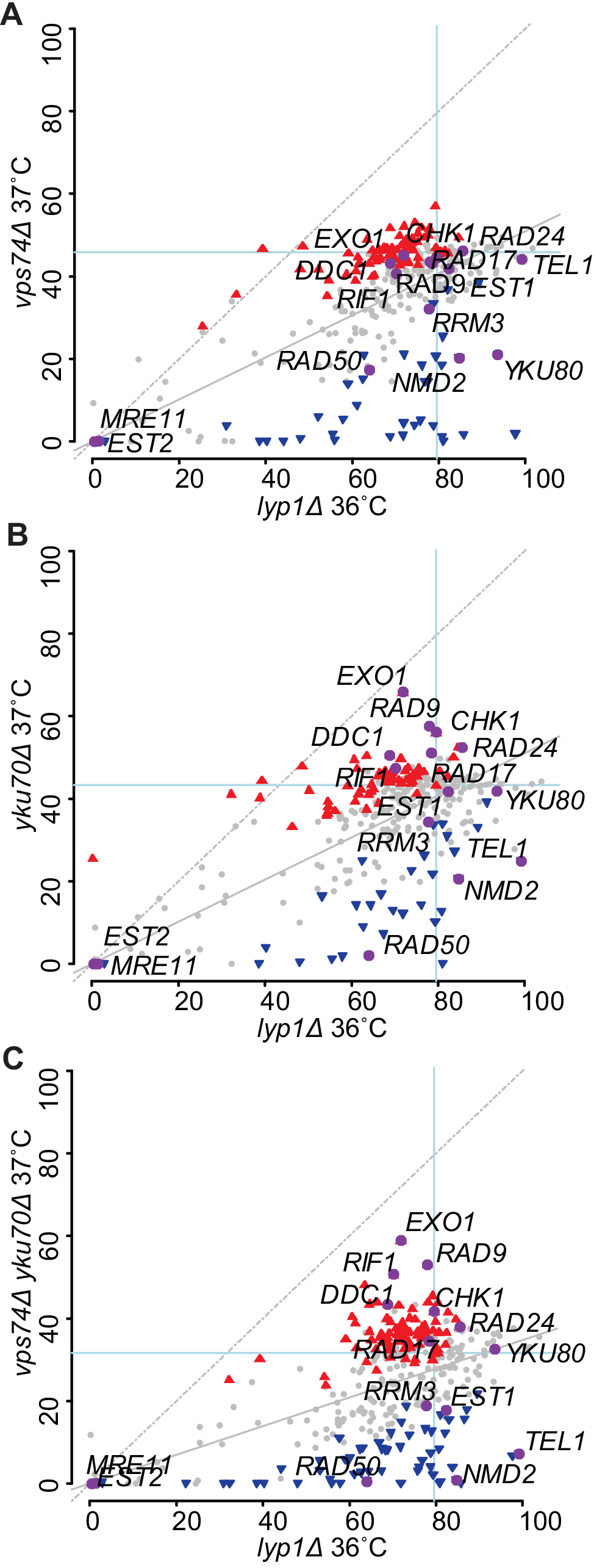
Systematic analysis of the effects of 358 gene deletions in the fitness of *vps74Δ, yku70Δ* and *yku70Δ vps74Δ* cells. Part of the yeast genome knock out collection (358 strains, Table S2) was crossed with *vps74Δ* (A), *yku70Δ* (B), *vps74Δ yku70Δ* (C) or *lyp1Δ* (A, B and C) mutations. Double or triple mutants were then grown on solid agar plates and the fitness was measured at 37°C (or 36°C for *lyp1Δ*). Fitness is measured as Maximum Doubling Rate × Maximum Doubling Potential (MDR × MDP, units are doublings squared per day, d2/day), as previously described (Addinall et al. 2011). Each dot indicates the effect of a gene deletion (*yfgΔ*) on the fitness of *vps74Δ* (A), *yku70Δ* (B), *vps74Δ yku70Δ* (C) or *lyp1Δ* (A, B and C). Grey dots indicate the deletions that did not significantly alter the fitness of the mutants in the *y* axis relative to the *x* axis (*lyp1Δ*). Blue inverted triangles represent gene deletions that are enhancers of the *vps74Δ* (A), *yku70Δ* (B) or *vps74Δ yku70Δ* (C), red triangles are suppressors and the purple dots represent the fitness of the deletion of 15 telomere related genes. Figures showing *vps74Δ* screens versus *yku70Δ* screens, *vps74Δ yku70Δ* screens versus *yku70Δ* screens and *vps74Δ yku70Δ* screens versus *vps74Δ* screens can be found in Figure S2B, C and D, respectively.

### Vps74 affects telomere biology similarly to Pmt1 and Pmt2

Our results show that Vps74 potentially collaborates with Yku70 to affect telomere function. However, since Vps74 is reported to be a cytoplasmic protein, its role at telomeres is likely to be indirect. GOLPH3, the mammalian orthologue of Vps74, is phosphorylated by DNA-PK upon DNA damage induction, and so it is possible that yeast Vps74 affects telomere biology through a similar phosphorylation pathway (Farber-Katz et al. 2014). In order to test if Vps74 is phosphorylated in response to different types of DNA damage, Vps74-13Myc was analysed by western blot after MMS treatment or in *cdc13-1* cells at 37°C. As a control, Rad53 was analysed as it is extensively phosphorylated upon DNA damage (Pellicioli et al. 1999; Morin et al. 2008). As previously reported, Rad53 was phosphorylated upon MMS treatment and in *cdc13-1* cells at 37°C, showing that the DNA damage response was activated in these cells (Figure S3). However, we found no evidence that Vps74 is phosphorylated in response to either type of DNA damage since we did not detect a mobility shift in Vps74-13Myc protein (Figure S3). Therefore, although we cannot exclude that Vps74 is phosphorylated in response to DNA damage we see no evidence that this is the case.

In yeast, Vps74 can be found in the cis and medial cisternae of the Golgi. Vps74 facilitates the function of mannosyltransferases, in glycosylation, and Sac1, a PtdIns4P phosphatase, to help modulate lipid levels and inositol phosphates, a class of intracellular signalling molecules (Schmitz et al. 2008; Wood et al. 2012; Short 2014) (Figure 5A). To try to understand if Vps74 affects *yku70Δ* cell fitness by affecting the function of mannosyltransferases or by affecting Sac1 function, we screened for yeast gene deletions that behave similarly to *vps74Δ* across five telomere related genetic screens (Figure 5B, Figure S4). To do this we used Profilyzer, a web based interaction tool (Dubarry et al. 2015; Holstein et al. 2017). Interestingly, *pmt1Δ* and *pmt2Δ*, affecting O-mannosyltransferases (transferring mannose from dolichyl phosphate-D-mannose to serine and threonine residues in proteins), behaved very similarly to *vsp74Δ* in *cdc13-1 exo1Δ*, *cdc13-1*, *stn1-13*, *yku70Δ* and *cdc13-1 rad9Δ* genetic screens (Figure 5B and Figure S4). On the other hand, *sac1Δ* caused extremely unfit cells in all screens (Figure 5B). The similarity between the effects of deleting *VPS74*, *PMT1* or *PMT2* in telomere defective cells, suggests that Vps74 affects the fitness of telomere defective cells through Pmt1 and Pmt2-dependent pathways. This supports the notion that Vps74 regulates telomere biology by affecting mannosyltransferase function. Interestingly, *mnn2Δ* (affecting an α-1,2-mannosyltransferase) also showed similar genetic interactions to *vps74Δ, pmt1Δ and pmt2Δ* across the telomere screens (Figure S4A-C).

**Figure 5.**
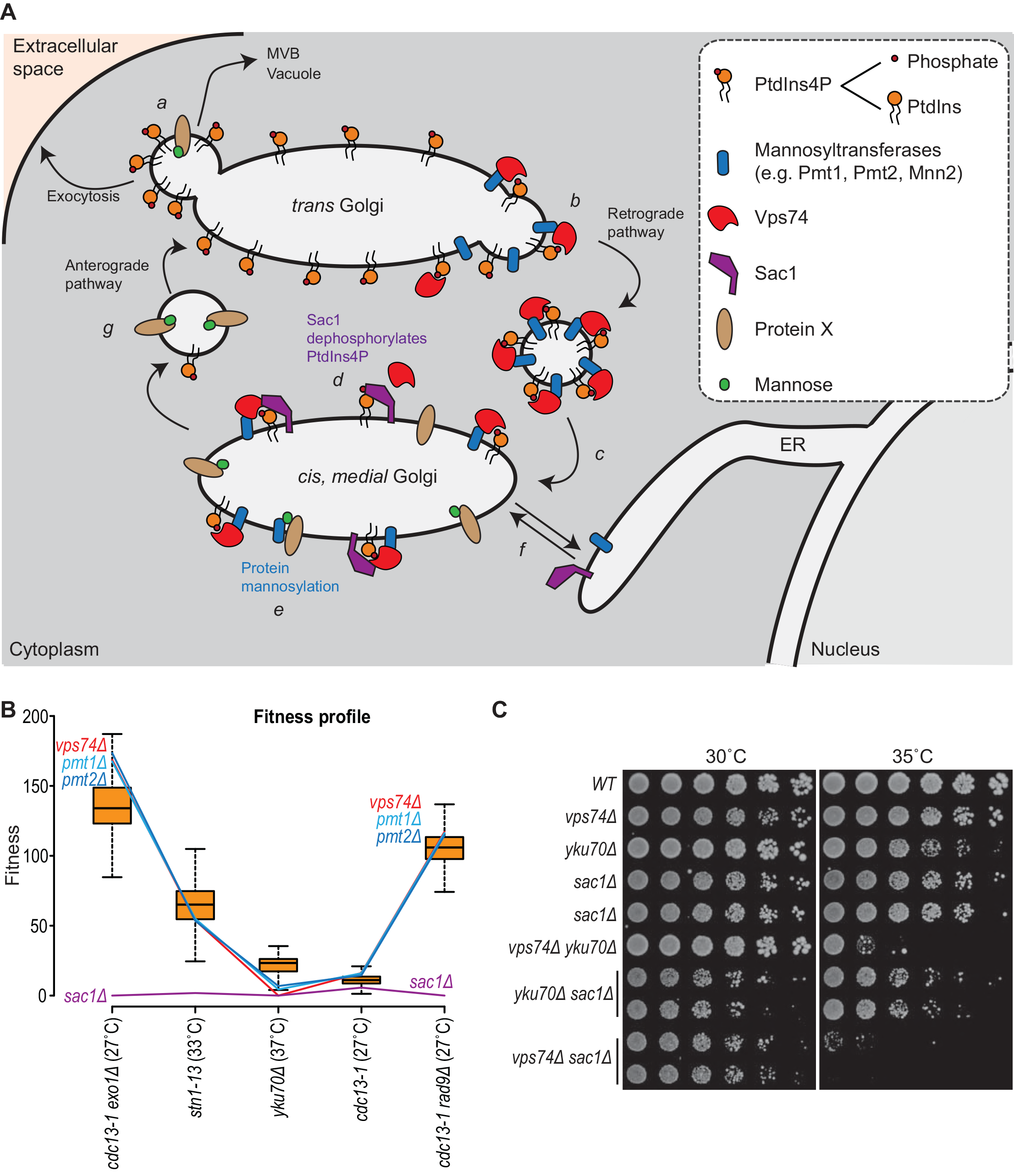
*vps74Δ*, *pmt1Δ* and *pmt2Δ* show similar genetic interactions with mutations affecting telomeres. **(A)** Cartoon showing the Vps74 function in protein sorting within the Golgi apparatus (Schmitz et al. 2008; Wood et al. 2012). Complex sphingolipids are preferentially packaged with secretory cargo into anterograde-directed transport vesicles and can end in exocytosis, multivesicular bodies (MVB) or the vacuole (a). In the trans Golgi, Vps74 recognizes PtdIns4P and mannosyltransferases, sorting them into COPI-coated retrograde vesicles (b). As a consequence of co-packaging PtdIns4P with Golgi protein residents into retrograde vesicles, PtdIns4P is delivered to the medial cisternae (c). In the medial/cis Golgi, Vps74 promotes Sac1-dependent PtdIns4P dephosphorylation (d) and protein mannosylation (e). Sac1 and mannosyltransferases cycle between the endoplasmic reticulum (ER) and the Golgi apparatus (f). Similarly, mannosylated proteins could follow the anterograde pathway (g, a) or eventually return to ER (f). Cartoon and legend adapted from (Wood et al. 2012). **(B)** Fitness profiles of *vps74Δ*, *sac1Δ*, *pmt1Δ* and *pmt2Δ* in combination with several mutations affecting telomere biology (*x* axis) (Holstein et al. 2017). Box plots show 50% range, the whiskers represent 1.5-fold the 50% range from the box, and the horizontal black line is the median fitness. **(C)** Spot test assays as described in Figure 1.

Although *sac1Δ* did not appear to behave similarly to *vps74Δ* across the telomere screens (Figure 5B), we decided to confirm this in a low-throughput manner in the W303 genetic background. *SAC1* was deleted in *vps74Δ*, *yku70Δ* and *vps74Δ yku70Δ* cells and fitness was measured. Interestingly, *vps74Δ* and *sac1Δ* are synthetically sick, even at 23°C, and this is not affected by*YKU70* (Figure 5C). This result suggests that Vps74 and Sac1 affect different pathways to maintain the fitness of yeast cells. Furthermore, *vps74Δ yku70Δ* cells are less fit than *sac1Δ yku70Δ* cells at 35°C, suggesting that Vps74 has a role in *yku70Δ* cells that is independent of Sac1. We conclude that Vps74 and Sac1 act in different ways to affect cell fitness of telomere proficient and deficient cells. Thus, the Vps74 role in telomere biology is at least partially independent of the *SAC1* pathway.

Finally, it cannot be excluded that Vps74 plays a more direct role in the nucleus, since Vps74-GFP localizes to the nucleus, as well as the cytoplasm (Figure S5). Indeed, using two nuclear localization prediction programs (cNLS mapper and NucPred), Vps74 is predicted to have a weak nuclear localization signal in its N-terminus (Brameier et al. 2007; Kosugi et al. 2009a; Kosugi et al. 2009b). We conclude that Vps74 regulates the fitness of *yku70Δ* cells independently of Sac1. Overall, genetic interaction data suggest that Vps74, Pmt1 and Pmt2 work in the same pathway to affect the fitness of telomere defective cells.

### Stn1 levels are regulated by a Vps74-dependent pathway

Since Vps74 is involved in at least two pathways likely to affect protein levels (protein glycosylation and PtdIns4P-dependent protein synthesis), we wondered if Vps74 might affect the levels of a protein or proteins that affect telomere biology. We noted that *nmd2Δ,* affecting nonsense mediated mRNA decay, causes similar phenotypes to *vps74Δ*, suppressing *cdc13-1* and enhancing *yku70Δ* fitness defects (Figure 1). *nmd2Δ,* like *vps74Δ,* also leads to telomere shortening (Figure 3C) (Addinall et al. 2011; Holstein et al. 2014). The effect of *nmd2Δ* has been ascribed to increased levels of Stn1 (Dahlseid et al. 2003), and indeed plasmid induced Stn1 overexpression suppressed *cdc13-1* and enhances *yku70Δ* fitness defects (Addinall et al. 2011). Stn1, along with Cdc13 and Ten1 are components of the CST complex involved in telomere capping (Addinall et al. 2011; Holstein et al. 2014). To test whether Vps74 affects Stn1 levels, Stn1-Myc levels were measured in *vps74Δ* cells. In addition, because *vps74Δ* cells show temperature dependent fitness defects (Figure 2), we measured Stn1 levels at 30°C and 37°C. Interestingly, Stn1 levels were increased approximately 40% in *vps74Δ* cells at 30°C (Figure 6A, B). Additionally, Stn1 levels were also increased by growth at 37°C. The increased Stn1 levels in *vps74Δ* cells could explain the short telomeres of these cells and the negative genetic interaction between *vps74Δ* and *yku70Δ* (Puglisi et al. 2008; Romano et al. 2013). It is interesting to speculate that increases in Stn1 levels at high temperature might help explain why telomeres get shorter in wild type cells at increased temperatures (Romano et al. 2013). We had observed that *vps74Δ* was synthetically sick with the *mre11Δ* mutation, and that *vps74Δ mre11Δ* double mutants have shorter telomeres than either single mutant (Figures 2D and 3). We wondered if the effect of *vps74Δ* in the *mre11Δ* context could be due to increased Stn1 levels. To test this we used a 2 micron plasmid to overexpress Stn1. Consistent with our hypothesis Stn1 overexpression reduced fitness of *mre11Δ* cells at all temperatures (Figure 6C). Overall we conclude that Vps74 helps maintain low levels of Stn1, and this may be the mechanism by which Vps74 affects telomere function in numerous different contexts.

**Figure 6.**
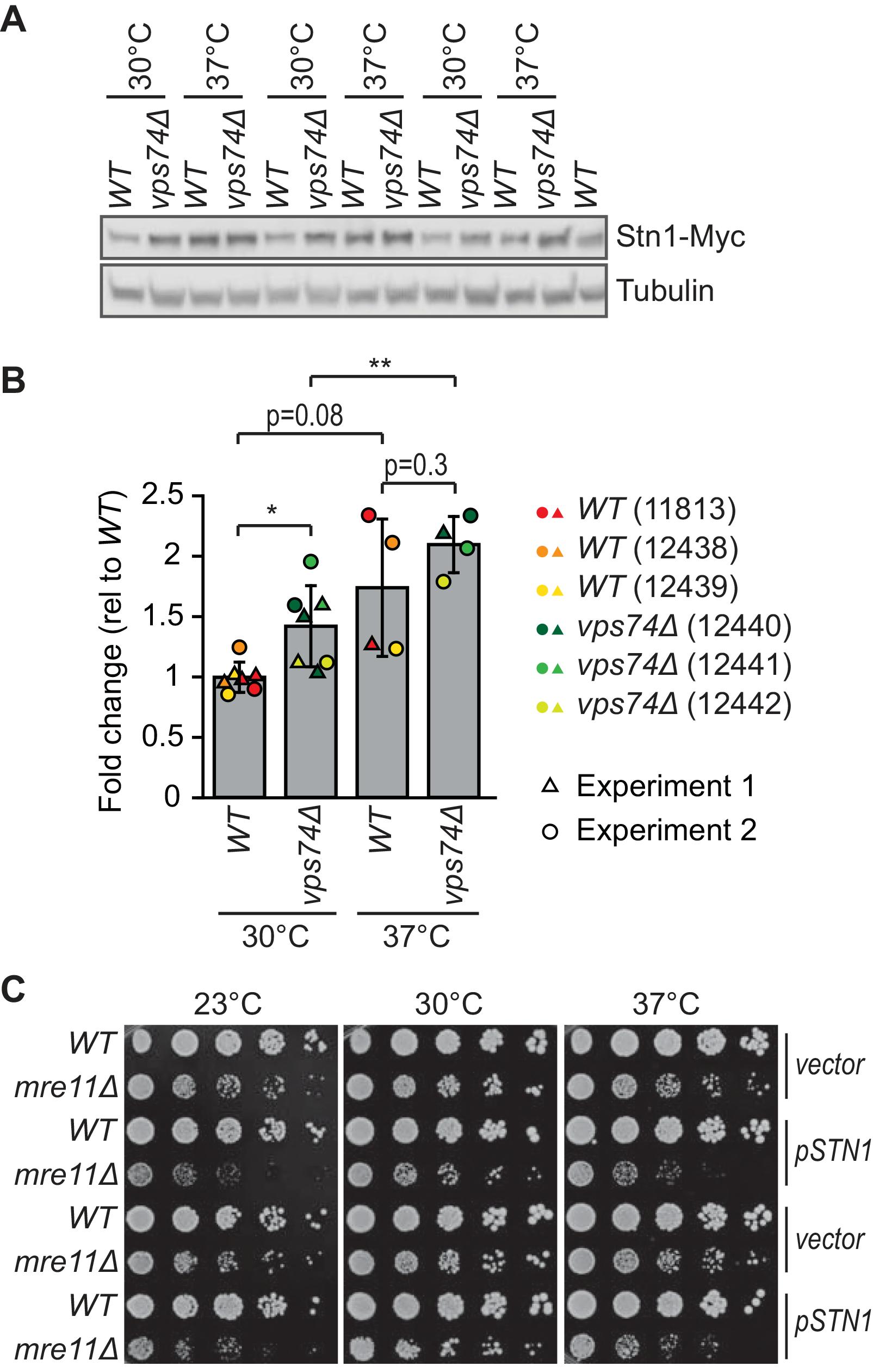
Increased temperature and loss of Vps74 leads to increased Stn1-Myc levels. Three independent *WT* and *vps74Δ* strains carrying a Stn1-Myc construct (strains numbers indicated), were cultured overnight at 30°C to saturation. Each culture was diluted 1:100, cultured for 2h at 30°C and then divided into two cultures that were further incubated at 30°C or 37°C for 4h. **(A)** Proteins were extracted using TCA and a western blot was performed first against Myc and then against tubulin. **(B)** Quantification of A and Figure S6 using Image J. Stn1-Myc intensity normalized for tubulin levels is shown. Values are presented as fold change relative to *WT* at 30°C. At 30°C, three independent strains were analysed in two independent experiments (n=6), while at 37° two independent strains were analysed once while a third independent strain was analysed twice (independent experiments, n=4). The mean is indicated and the error bars indicate the standard deviation. Statistical analyses used the two-tailed t-Test assuming unequal variance (*P<0.05 and **P<0.01) performed with SigmaPlot (version 11). **(C)** *WT* and *mre11Δ* cells were transformed with a 2 μm plasmid carrying *STN1* or a vector plasmid (plasmids described in Table S3). Six independent (*WT* and *mre11Δ*) transformants carrying either the *STN1* plasmid or the vector, were cultivated in selective media (-LEU, lacking leucine) until saturation and spot tests were performed as described in Figure 1 (in −LEU plates). Pictures were taken after 3 days of incubation. Two representative strains of each genotype carrying either of the plasmids are shown.

## DISCUSSION

In mammalian cells, GOLPH3 phosphorylation by DNA-PK was shown to be essential for Golgi fragmentation in response to DNA damage (Farber-Katz et al. 2014). The functional implications of Golgi fragmentation are not yet clear in mammalian cells. Interestingly, yeast genetic data also support the existence of a connection between Golgi and the DNA damage response (DDR), indicating that this relationship might be evolutionary conserved. Specifically, yeast genome-wide genetic interaction screens suggested that the yeast ortholog of *GOLPH3*, *VPS74,* affected the fitness of telomere defective *cdc13-1* and *yku70Δ* cells (Addinall et al. 2011). There is an intimate and complex relationship between functional telomeres and the DDR, with many DDR proteins affecting telomere function and vice versa (Lydall 2009).

In agreement with the published high-throughput data, our new low-throughput experiments showed that *vps74Δ* suppressed *cdc13-1* fitness defects and enhanced *yku70Δ* defects (Addinall et al. 2011). These data confirm that yeast Vps74 affects telomere function and are consistent with a role for Vps74 in the DNA damage response. Our evidence suggests that the better fitness of *vps74Δ cdc13-1* cells in comparison to *cdc13-1* cells is due to a decrease in the telomeric ssDNA levels in the double mutants. On the other hand, the decrease in the fitness of *vps74Δ yku70Δ* cells seems to be independent of telomeric ssDNA. Interestingly, numerous gene deletions that suppressed or enhanced *yku70Δ* temperature defects, similarly affected the temperature sensitivity of *vps74Δ* and *vps74Δ yku70Δ* cells, suggesting that Vps74 and Yku70 perform similar functions at telomeres. Amongst the confirmed strong suppressors of *vps74Δ yku70Δ* temperature defects were *exo1Δ*, *chk1Δ*, *mec1Δ* (*sml1Δ*), affecting DDR genes, that also suppress *yku70Δ* fitness defects (Maringele and Lydall 2002). Thus, our genetic interaction data suggests that Vps74 contributes in some manner to telomere capping, and this role is more important in the absence of Yku70.

It has previously been reported that *nmd2Δ*, affecting a core component of the nonsense-mediated mRNA decay pathway, improved the fitness of *cdc13-1* cells and decreased the fitness of *yku70Δ* cells by increasing levels of Stn1 (Dahlseid et al. 2003; Addinall et al. 2011; Holstein et al. 2014). Interestingly, *nmd2Δ* and *vps74Δ* genetic interaction patterns are similar in *cdc13-1* and *yku70Δ* contexts. Stn1 is an essential telomere capping protein and functions with Cdc13 and Ten1 in the CST complex. Increased levels of Stn1 are thought to help a partially defective Cdc13-1 protein cap the chromosome end (Holstein et al. 2014). On the other hand, high levels of Stn1 inhibit Cdc13-dependent recruitment of telomerase to telomeres, causing a short telomere phenotype (Grandin et al. 2000). Stn1 overexpression enhances growth defects of *yku70Δ* mutants with short telomeres, presumably by exacerbating the short telomere phenotype (Dahlseid et al. 2003; Addinall et al. 2011; Holstein et al. 2014). Interestingly, in *vps74Δ* cells, as in *nmd2Δ* cells, increased levels of Stn1, can explain the suppression of *cdc13-1* and enhancement *yku70Δ* cell fitness defects. Importantly, increased Stn1 levels could also explain the short telomeres of *vps74Δ* cells and the poor fitness of *vps74Δ mre11Δ* cells, since *mre11Δ* cells, like *yku70Δ* cells, have a short telomere phenotype. For all these reasons we propose that Vps74 affects telomere function by maintaining low levels of Stn1.

Vps74 regulation of Stn1 levels is unlikely to be direct since all known Vps74 functions are in the Golgi and Stn1 is a nuclear protein (Schmitz et al. 2008; Wood et al. 2012; Cai et al. 2014; Short 2014). High-throughput genetic interactions suggest that *VPS74*, *PMT1* and *PMT2* could work in the same pathways to affect telomere function. Pmt1 and Pmt2 are responsible for O-mannosylation of membrane proteins, affecting the stability/function of such proteins (Schmitz et al. 2008; Petkova et al. 2012; Loibl and Strahl 2013; Short 2014). We speculate that Vps74 (and Pmt1/Pmt2) affect the levels of nuclear proteins, like Stn1, by affecting signal transduction pathways whose cell surface components are targets of mannosyltransferases. For example, we suggest that Vps74 may be important for the Golgi localization and function of Pmt1 and Pmt2 (Figure S7, a) (Schmitz et al. 2008). It is known that Pmt1 and Pmt2 are responsible for the mannosylation of Mtl1, a cytoplasmic transmembrane sensor protein upstream of the cytoplasmic Pkc1-MAPK pathway (Figure S7, b) (Petkova et al. 2012). Pck1 activates Bck1, which in turn activates Mkk1/2, which finally activates Mpk1, involved in nuclear and cytoplasmic responses to oxidative and genotoxic stresses, including transcriptional modulation and proteasome homeostasis (Figure S7, c) (Truman et al. 2009; Jendretzki et al. 2011; Soriano-Carot et al. 2012; Rousseau and Bertolotti 2016). Thus, it seems likely that Vps74 affects the levels of nuclear proteins, such as Stn1, by modulating signal transduction pathways that depend on protein mannosylation.

In human cancer cells, GOLPH3 overexpression was associated with increased activation of mTOR signalling, affecting protein synthesis in response to nutrient changes (Scott et al. 2009). Perhaps, therefore, Vps74, Pmt1 and Pmt2 affect the levels of Stn1 (and likely other proteins) by affecting the Tor and Pkc1-MAPK pathways (Figure S7). Overall, our careful analysis of Vps74 function in telomere defective yeast cells, leads us to believe that regulation of Stn1 levels is at least one of the ways Vps74 affects telomere function. Future experiments will be required to better understand the mechanisms by which Vps74 affects Stn1 levels, the DNA damage response and telomere functions. It will also be interesting to determine if these mechanisms are conserved across eukaryotes.

## REFERENCES

Adams A, Gottschcling DE, Kaiser CA, Stearns T. 1997. Methods in Yeast Genetics:. Cold Spring Harbor Laboratory Press, New York.

Addinall SG, Holstein EM, Lawless C, Yu M, Chapman K, Banks AP, Ngo HP, Maringele L, Taschuk M, Young A et al. 2011. Quantitative Fitness Analysis Shows That NMD Proteins and Many Other Protein Complexes Suppress or Enhance Distinct Telomere Cap Defects. PLoS genetics 7.

Brameier M, Krings A, MacCallum RM. 2007. NucPred–predicting nuclear localization of proteins. Bioinformatics 23: 1159–1160.

Cai Y, Deng Y, Horenkamp F, Reinisch KM, Burd CG. 2014. Sac1-Vps74 structure reveals a mechanism to terminate phosphoinositide signaling in the Golgi apparatus. The Journal of cell biology 206: 485–491.

Dahlseid JN, Lew-Smith J, Lelivelt MJ, Enomoto S, Ford A, Desruisseaux M, McClellan M, Lue N, Culbertson MR, Berman J. 2003. mRNAs encoding telomerase components and regulators are controlled by UPF genes in Saccharomyces cerevisiae. Eukaryotic cell 2: 134–142.

Dewar JM, Lydall D. 2010. Pif1- and Exo1-dependent nucleases coordinate checkpoint activation following telomere uncapping. Embo Journal 29: 4020–4034.

Dippold HC, Ng MM, Farber-Katz SE, Lee SK, Kerr ML, Peterman MC, Sim R, Wiharto PA, Galbraith KA, Madhavarapu S et al. 2009. GOLPH3 bridges phosphatidylinositol-4-phosphate and actomyosin to stretch and shape the Golgi to promote budding. Cell 139: 337–351.

Dubarry M, Lawless C, Banks AP, Cockell S, Lydall D. 2015. Genetic Networks Required to Coordinate Chromosome Replication by DNA Polymerases alpha, delta, and epsilon in Saccharomyces cerevisiae. G3 5: 2187–2197.

Farber-Katz SE, Dippold HC, Buschman MD, Peterman MC, Xing M, Noakes CJ, Tat J, Ng MM, Rahajeng J, Cowan DM et al. 2014. DNA damage triggers Golgi dispersal via DNA-PK and GOLPH3. Cell 156: 413–427.

Glick BS, Nakano A. 2009. Membrane traffic within the Golgi apparatus. Annual review of cell and developmental biology 25: 113–132.

Goldstein AL, McCusker JH. 1999. Three new dominant drug resistance cassettes for gene disruption in Saccharomyces cerevisiae. Yeast 15: 1541–1553.

Grandin N, Damon C, Charbonneau M. 2000. Cdc13 cooperates with the yeast Ku proteins and Stn1 to regulate telomerase recruitment. Molecular and cellular biology 20: 8397–8408.

Holstein EM, Clark KR, Lydall D. 2014. Interplay between nonsense-mediated mRNA decay and DNA damage response pathways reveals that Stn1 and Ten1 are the key CST telomere-cap components. Cell Rep 7: 1259–1269.

Holstein EM, Ngo G, Lawless C, Banks P, Greetham M, Wilkinson D, Lydall D. 2017. Systematic Analysis of the DNA Damage Response Network in Telomere Defective Budding Yeast. G3 7: 2375–2389.

Jendretzki A, Wittland J, Wilk S, Straede A, Heinisch JJ. 2011. How do I begin? Sensing extracellular stress to maintain yeast cell wall integrity. Eur J Cell Biol 90: 740–744.

Kosugi S, Hasebe M, Matsumura N, Takashima H, Miyamoto-Sato E, Tomita M, Yanagawa H. 2009a. Six classes of nuclear localization signals specific to different binding grooves of importin alpha. The Journal of biological chemistry 284: 478–485.

Kosugi S, Hasebe M, Tomita M, Yanagawa H. 2009b. Systematic identification of cell cycle-dependent yeast nucleocytoplasmic shuttling proteins by prediction of composite motifs. Proc Natl Acad Sci U S A 106: 10171–10176.

Loibl M, Strahl S. 2013. Protein O-mannosylation: What we have learned from baker’s yeast. Bba-Mol Cell Res 1833: 2438–2446.

Lydall D. 2009. Taming the tiger by the tail: modulation of DNA damage responses by telomeres. The EMBO journal 28: 2174–2187.

Maringele L, Lydall D. 2002. EXO1-dependent single-stranded DNA at telomeres activates subsets of DNA damage and spindle checkpoint pathways in budding yeast yku70 Delta mutants. Genes Dev 16: 1919–1933.

Maringele L, Lydall D. 2004. EXO1 plays a role in generating type I and type II survivors in budding yeast. Genetics 166: 1641–1649.

Morin I, Ngo HP, Greenall A, Zubko MK, Morrice N, Lydall D. 2008. Checkpoint-dependent phosphorylation of Exo1 modulates the DNA damage response. The EMBO journal 27: 2400–2410.

Ngo GH, Lydall D. 2015. The 9-1-1 checkpoint clamp coordinates resection at DNA double strand breaks. Nucleic acids research 43: 5017–5032.

Pellicioli A, Lucca C, Liberi G, Marini F, Lopes M, Plevani P, Romano A, Di Fiore PP, Foiani M. 1999. Activation of Rad53 kinase in response to DNA damage and its effect in modulating phosphorylation of the lagging strand DNA polymerase. The EMBO journal 18: 6561–6572.

Petkova MI, Pujol-Carrion N, de la Torre-Ruiz MA. 2012. Mtl1 O-mannosylation mediated by both Pmt1 and Pmt2 is important for cell survival under oxidative conditions and TOR blockade. Fungal Genet Biol 49: 903–914.

Potelle S, Klein A, Foulquier F. 2015. Golgi post-translational modifications and associated diseases. J Inherit Metab Dis 38: 741–751.

Puglisi A, Bianchi A, Lemmens L, Damay P, Shore D. 2008. Distinct roles for yeast Stn1 in telomere capping and telomerase inhibition. The EMBO journal 27: 2328–2339.

Romano GH, Harari Y, Yehuda T, Podhorzer A, Rubinstein L, Shamir R, Gottlieb A, Silberberg Y, Pe’er D, Ruppin E et al. 2013. Environmental stresses disrupt telomere length homeostasis. PLoS genetics 9: e1003721.

Rousseau A, Bertolotti A. 2016. An evolutionarily conserved pathway controls proteasome homeostasis. Nature 536: 184–189.

Schmitz KR, Liu J, Li S, Setty TG, Wood CS, Burd CG, Ferguson KM. 2008. Golgi localization of glycosyltransferases requires a Vps74p oligomer. Dev Cell 14: 523–534.

Scott KL, Kabbarah O, Liang MC, Ivanova E, Anagnostou V, Wu J, Dhakal S, Wu M, Chen S, Feinberg T et al. 2009. GOLPH3 modulates mTOR signalling and rapamycin sensitivity in cancer. Nature 459: 1085–1090.

Shental-Bechor D, Levy Y. 2008. Effect of glycosylation on protein folding: a close look at thermodynamic stabilization. Proc Natl Acad Sci U S A 105: 8256–8261.

Short B. 2014. Vps74 gives phosphatase directions. Journal of Cell Biology 206: 453–453.

Soriano-Carot M, Bano MC, Igual JC. 2012. The yeast mitogen-activated protein kinase Slt2 is involved in the cellular response to genotoxic stress. Cell Div 7: 1.

Strahl T, Thorner J. 2007. Synthesis and function of membrane phosphoinositides in budding yeast, Saccharomyces cerevisiae. Biochimica et biophysica acta 1771: 353–404.

Tong AH, Boone C. 2006. Synthetic genetic array analysis in Saccharomyces cerevisiae. Methods in molecular biology 313: 171–192.

Tong AHY, Evangelista M, Parsons AB, Xu H, Bader GD, Page N, Robinson M, Raghibizadeh S, Hogue CWV, Bussey H et al. 2001. Systematic genetic analysis with ordered arrays of yeast deletion mutants. Science 294: 2364–2368.

Truman AW, Kim KY, Levin DE. 2009. Mechanism of Mpk1 mitogen-activated protein kinase binding to the Swi4 transcription factor and its regulation by a novel caffeine-induced phosphorylation. Molecular and cellular biology 29: 6449–6461.

Wera S, Bergsma JC, Thevelein JM. 2001. Phosphoinositides in yeast: genetically tractable signalling. FEMS yeast research 1: 9–13.

Wood CS, Hung CS, Huoh YS, Mousley CJ, Stefan CJ, Bankaitis V, Ferguson KM, Burd CG. 2012. Local control of phosphatidylinositol 4-phosphate signaling in the Golgi apparatus by Vps74 and Sac1 phosphoinositide phosphatase. Molecular biology of the cell 23: 2527–2536.

Xu C, Ng DT. 2015. Glycosylation-directed quality control of protein folding. Nature reviews Molecular cell biology 16: 742–752.

